# Bile acid-sensitive human norovirus strains are susceptible to sphingosine-1-phosphate receptor 2 inhibition

**DOI:** 10.1101/2024.01.02.573926

**Authors:** Victoria Tenge, B. Vijayalakshmi Ayyar, Khalil Ettayebi, Sue E. Crawford, Yi-Ting Shen, Frederick H. Neill, Robert L. Atmar, Mary K. Estes

## Abstract

Human noroviruses (HuNoVs) are a diverse group of RNA viruses that cause both endemic and pandemic acute viral gastroenteritis. Previously we reported that many strains of HuNoV require bile or bile acid (BA) to infect human jejunal intestinal enteroid cultures. Of note, BA was not essential for replication of a pandemic-causing GII.4 HuNoV strain. Using the BA-requiring strain GII.3, we found that the hydrophobic BA GCDCA induces multiple cellular responses that promote replication in jejunal enteroids. Further, we found that chemical inhibition of the G-protein coupled receptor, sphingosine-1- phosphate receptor 2 (S1PR2), by JTE-013 reduced both GII.3 infection in a dose-dependent manner and cellular uptake in enteroids. Herein, we sought to determine if S1PR2 is required by other BA-dependent HuNoV strains and BA-independent GII.4, and if S1PR2 is required for BA-dependent HuNoV infection in other segments of the small intestine. We found JTE-013 inhibition of S1PR2 in jejunal HIEs reduces GI.1, GII.3, and GII.17 (BA-dependent) but not the GII.4 Sydney variant (BA-independent) infection, providing additional evidence of strain-specific differences in HuNoV infection. GII.3 infection of duodenal, jejunal and ileal lines derived from the same individual was also reduced with S1PR2 inhibition, indicating a common mechanism of BA-dependent infection among multiple segments of the small intestine. Our results support a model where BA-dependent HuNoV exploit the activation of S1PR2 by BA to infect the entire small intestine.

**Importance:** Human noroviruses (HuNoVs) are important viral human pathogens that cause both outbreaks and sporadic gastroenteritis. These viruses are diverse, and many strains are capable of infecting humans. Our previous studies have identified strain-specific requirements for hydrophobic bile acids (BAs) to infect intestinal epithelial cells. Moreover, we identified a BA receptor, sphingosine-1-phosphate receptor 2 (S1PR2), required for infection by a BA-dependent strain. To better understand how various HuNoV strains enter and infect the small intestine and the role of S1PR2 in HuNoV infection, we evaluated infection by additional HuNoV strains using an expanded repertoire of intestinal enteroid cell lines. We found that multiple BA-dependent strains, but not a BA- independent strain, all required S1PR2 for infection. Additionally, BA-dependent infection required S1PR2 in multiple segments of the small intestine. Together these results indicate S1PR2 has value as a potential therapeutic target for BA-dependent HuNoV infection.

## Introduction

HuNoVs are members of the *Caliciviridae*, a family of single-stranded, positive sense RNA viruses and are a leading cause of viral gastroenteritis worldwide. These viruses cause outbreaks, sporadic disease, and foodborne illness. Symptoms of HuNoV-induced disease include vomiting and diarrhea, and adverse outcomes are more likely to occur in young children, the immunocompromised, and the elderly (1–3). Since the introduction of the rotavirus vaccine, HuNoV has replaced rotavirus as the leading cause of viral gastroenteritis in children under the age of 5 years (4). Some immunocompromised individuals can develop chronic HuNoV infection (5). Globally, these viruses cause over 200,000 deaths with an economic impact of more than $60 billion in direct and indirect costs annually (6, 7). In many cases, HuNoV infection can be managed through oral rehydration to prevent dehydration. Antiemetics and antimotility drugs can be administered to reduce vomiting or diarrhea, but neither of these therapies are appropriate for use in children. For those individuals with severe or persistent disease, there is a significant need for development of therapies. Unfortunately, there are no licensed antivirals or therapeutics to combat this important group of viruses.

One challenge to identification of therapeutic targets is that HuNoVs are genetically diverse. Currently, there are 10 designated and 2 tentative genogroups (GI-GX, GNA1, GNA2), of which GI, GII, GIV, GVIII, and GIX contain viruses that infect humans (8). GI contains genotypes GI.1-GI.9 and GII contains GII.1-GII.27 and two tentative genotypes (8). Recent development of human intestinal enteroids (HIEs) for HuNoV cultivation has allowed us to begin exploring replication of many different genotypes of HuNoV and identify strain-specific requirements for infection (9–14). One such difference is the requirement for BAs in infection. The pandemic-causing GII.4 Sydney strain can replicate in HIEs independently of BA addition and enters HIEs by using wound repair and clathrin-independent carrier pathways (14), whereas another strain (e.g. GII.3) requires the addition of bile or hydrophobic BA for viral entry and infection (11). Unexpectedly, we found that the hydrophobic BA, GCDCA, triggers endosomal uptake required for GII.3 infection of a jejunal HIE line by the G-protein coupled receptor sphingosine-1-phosphate receptor 2 (S1PR2) (11). The established ligand for S1PR2 is the lipid sphingosine-1-P (S1P), but recently BAs were demonstrated to act on S1PR2 in both the liver and the intestine (15–17). Inhibition of S1PR2 activity by the chemical JTE-013 blocks both GII.3 uptake and infection in a dose-dependent manner (11). Seeking to expand these studies, we used the HIE HuNoV infection system to explore the effect of S1PR2 inhibition on replication of BA-independent GII.4 and two additional important BA-dependent HuNoV strains (GII.17 and GI.1). We also used infections of jejunal HIE lines derived from several patients as well as matched duodenal, jejunal and ileal lines derived from two individuals.

In the current study, we show that while GII.4 infection is not affected by inhibition of S1PR2, the replication of BA-dependent GII.3, GII.17 and GI.1 strains is reduced in several jejunal HIE lines upon S1PR2 inhibition, as is the appearance of BA-induced apical ceramide. GII.3 infection requires S1PR2 in all segments of the small intestine (duodenum, jejunum, and ileum) for replication. BA treatment also increases detection of S1PR2 protein, establishing a direct association between S1PR2 and replication of BA-dependent strains. Combined, these results indicate the existence of distinct entry and infection requirements for BA-independent versus BA-dependent strains and highlight the potential for therapeutically targeting BA-dependent HuNoV strains by S1PR2 inhibition.

## Results

### GII.3 infection of multiple jejunal lines is reduced in the presence of JTE-013

In our initial studies on GII.3 entry mechanisms and requirement for the S1PR2 receptor, we worked primarily with a secretor-positive jejunal HIE line, J2 (11). Subsequently, we transitioned to the use of commercial IntestiCult™ Organoid Growth Medium (OGM) for plating and differentiating monolayers, which has enhanced HuNoV replication in HIEs (13). To confirm our initial findings, we repeated inhibition experiments with J2 as a control and a panel of susceptible secretor-positive HIE lines with different genetic backgrounds grown in OGM. Among these lines is J4FUT2KI, a genetically modified line. The J4 HIE line is genetically and phenotypically secretor-negative and not susceptible to infection by many HuNoV strains (10). We modified this line by overexpression of FUT2 to generate a secretor-positive HuNoV susceptible line (10). Infection levels in J4FUT2KI HIEs are generally higher than observed in infection of J2, making this line suitable for testing drug inhibition of less robustly infecting strains (10, 18). We infected all lines in the presence of the S1PR2 inhibitor JTE-013 or vehicle control and, as expected, GII.3 infection of J2 in the presence of JTE-013 was reduced in comparison to the control (**Figure 1A**). In line with these results, GII.3 infection of other jejunal lines J3 and J11 was also inhibited when JTE-013 was present (**Figure 1A**). Replication of GII.3 in the genetically modified line, J4FUT2KI, was similarly reduced in the presence of JTE-013, indicating infection of this line behaves similarly to the unmodified HIE lines in response to chemical inhibition (**Figure 1B**).

**Figure 1.**
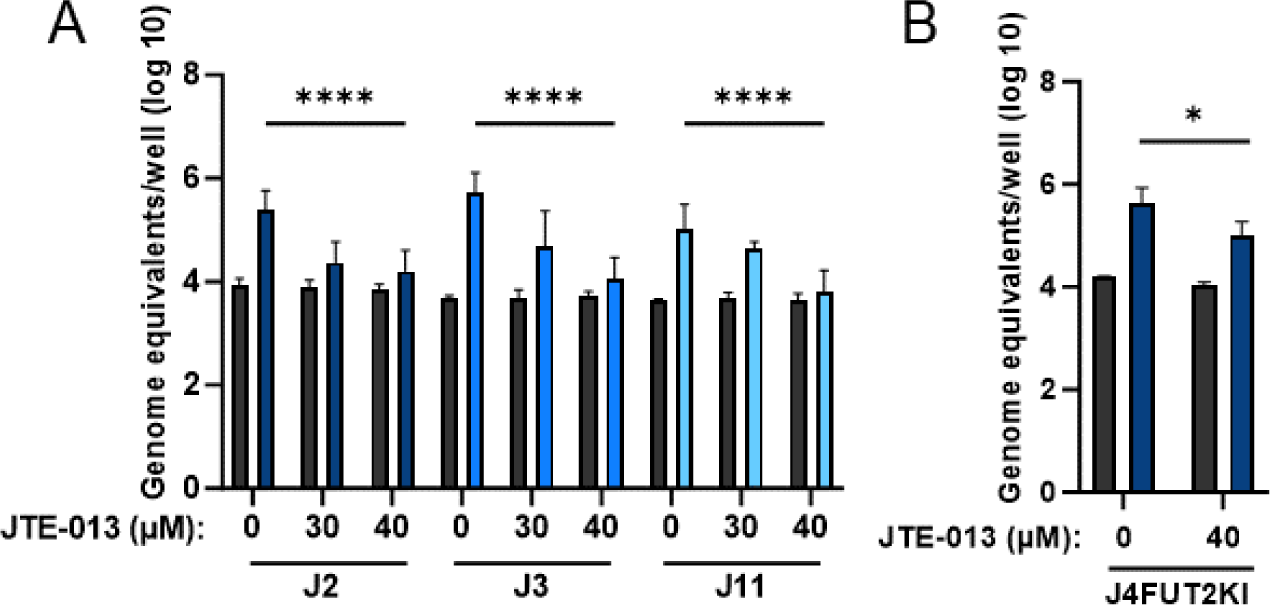
GII.3 infection of multiple jejunal lines is reduced in the presence of JTE-013. GII.3 (4.3 × 105 GEs/well) virus replication was evaluated in four jejunal (A: J2, J3, and J11; B: J4FUT2KI) HIE lines plated in OGM. Each experiment was performed twice, and compiled data are presented. Error bars denote standard deviation (n = 12). Significance was determined using two-way ANOVA (P value -*, < 0.05; ****, 0.001).

### BA treatment increases detection of S1PR2 in HIEs and inhibition of S1PR2 reduces BA-induced apical ceramide appearance

We previously reported a transient increase in GCDCA-induced endosomal acidification and subsequent endocytosis in HIEs 10 minutes after treatment (11). This led us to investigate expression of the BA receptor S1PR2 in J2 HIEs incubated with GCDCA for a short time-period (10 minutes). Western blotting of J2 cell lysates and confocal analysis of cells with or without GCDCA treatment showed that the presence of GCDCA significantly increases detection of expressed S1PR2 (**Figure 2**).

**Figure 2.**
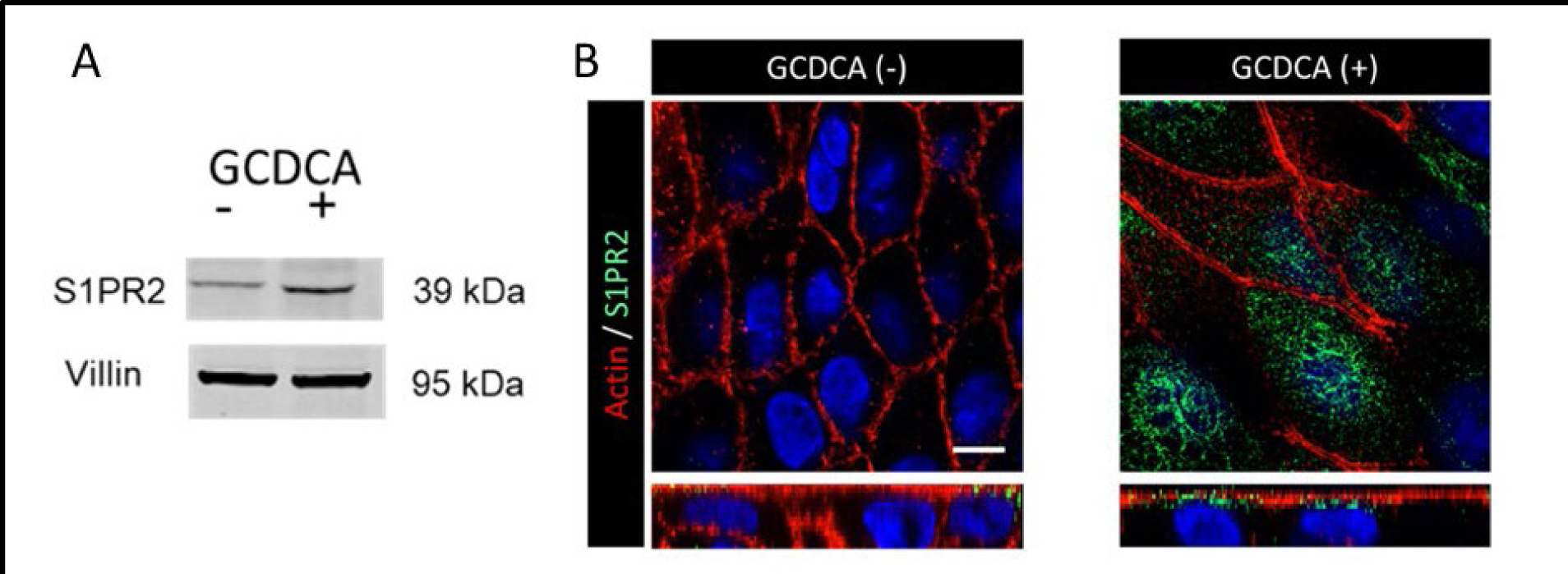
Exogeneous addition of GCDCA increases detection of S1PR2 in HIEs. A. Western blotting of J2 HIE lysates from cells without and with GCDCA (500 μM) treatment for 10 min at 37°C. Quantitation of S1PR2, relative to the villin loading control, shows a 2.58 increase in S1PR2 expression in cells treated with GCDCA, n=3. **B.** Confocal microscopy shows detection of S1PR2 (green) in J2 HIEs treated without and with GCDCA (500 μM). Actin (red) was detected using phalloidin. Top panels show 2D views whereas the bottom panels show orthogonal views of the HIEs. Bar, 5 μm.

We previously showed that BA also induces the appearance of ceramide on the apical surface of HIEs as well as exocytosis of LAMP-1 onto the cell surface of HIEs (11). Using JTE-013 treatment, we found S1PR2 was required for each of these rapid molecular changes (**Figure 3**). These data confirm the importance of S1PR2 in the BA- dependent events previously shown to be required for GII.3 infection.

**Figure 3.**
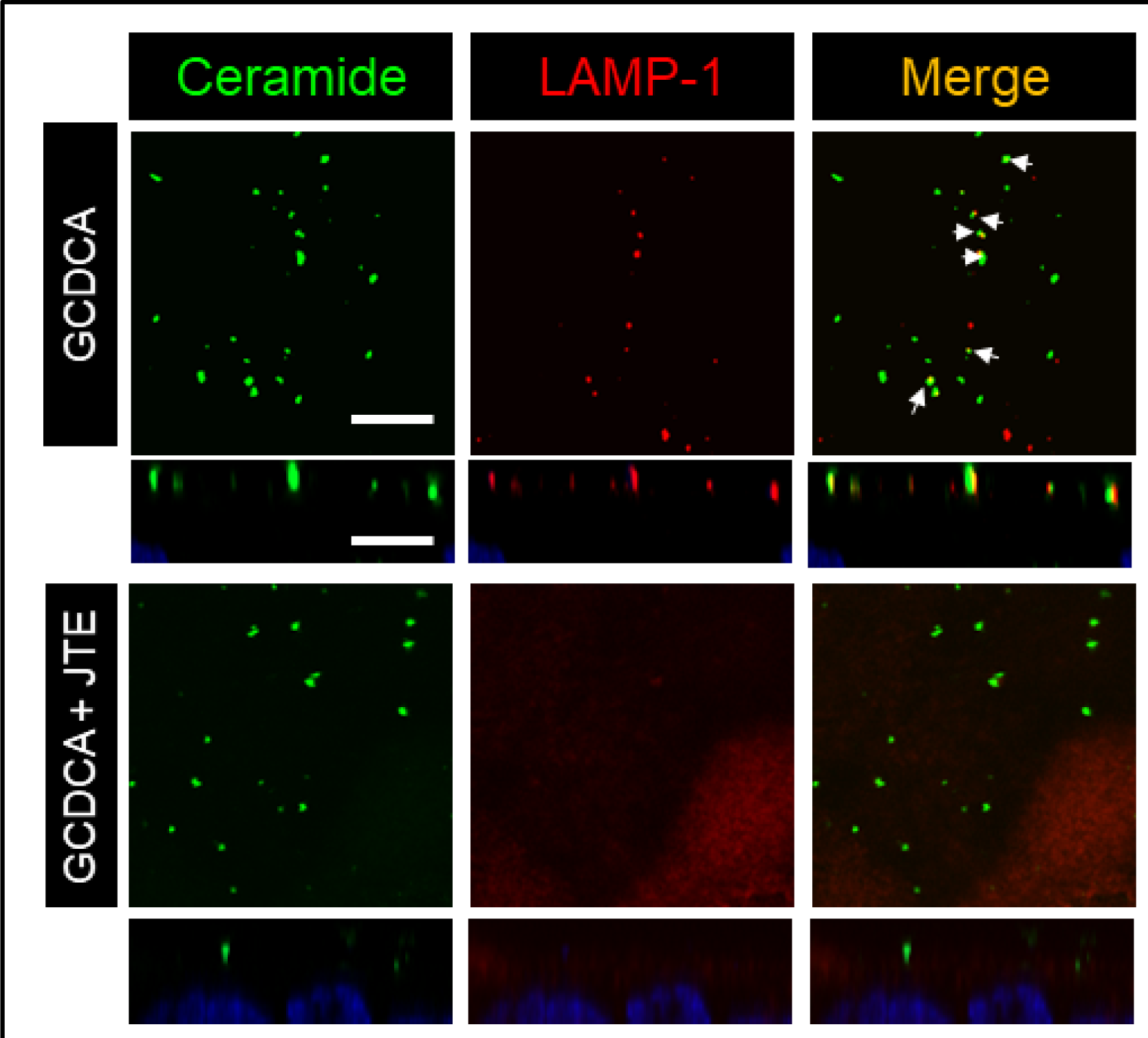
Inhibition of S1PR2 reduces GCDCA-induced apical ceramide appearance and LAMP-1 exocytosis onto the cell surface of HIEs. Confocal microscopy shows detection of ceramide (green) and LAMP-1 (red) in J2 HIEs. The HIEs were treated with GCDCA (500 μM, top panel) at 37oC for 10 min in the absence and presence of the S1PR2 inhibitor, JTE-013. White arrows represent ceramide and LAMP-1 colocalization in JTE-013-treated vs untreated cells. Top panels show 2D views, whereas the bottom panels show orthogonal views of the HIEs. Bar, 5 μm.

### BA-independent HuNoV infection is not affected by JTE-013

GII.3 and GII.4 HuNoV strains have strain-specific differences in infection, including whether they require BA for entry (11). Despite this, both viruses share the requirement for acid sphingomyelinase activity, an enzyme required for ceramide production from the lipid sphingomyelin, indicating they might have some shared entry requirements (11, 14). To determine if S1PR2 is required for BA-independent GII.4 infection, we infected J2 HIEs in the presence of JTE-013 using the Sydney/2012 variant. Moreover, because enhanced infection of GII.4 in the presence of bile has been observed (9), we also performed this experiment in the presence of GCDCA. Infection with 500 µM GCDCA was greater than but not statistically different from infection without GCDCA, and the presence of JTE-013 had no effect on infection whether or not GCDCA was added to the media (**Figure 4**). This is further evidence of differences in requirements between the pandemic causing, BA-independent GII.4s and BA-dependent strains of HuNoV.

**Figure 4.**
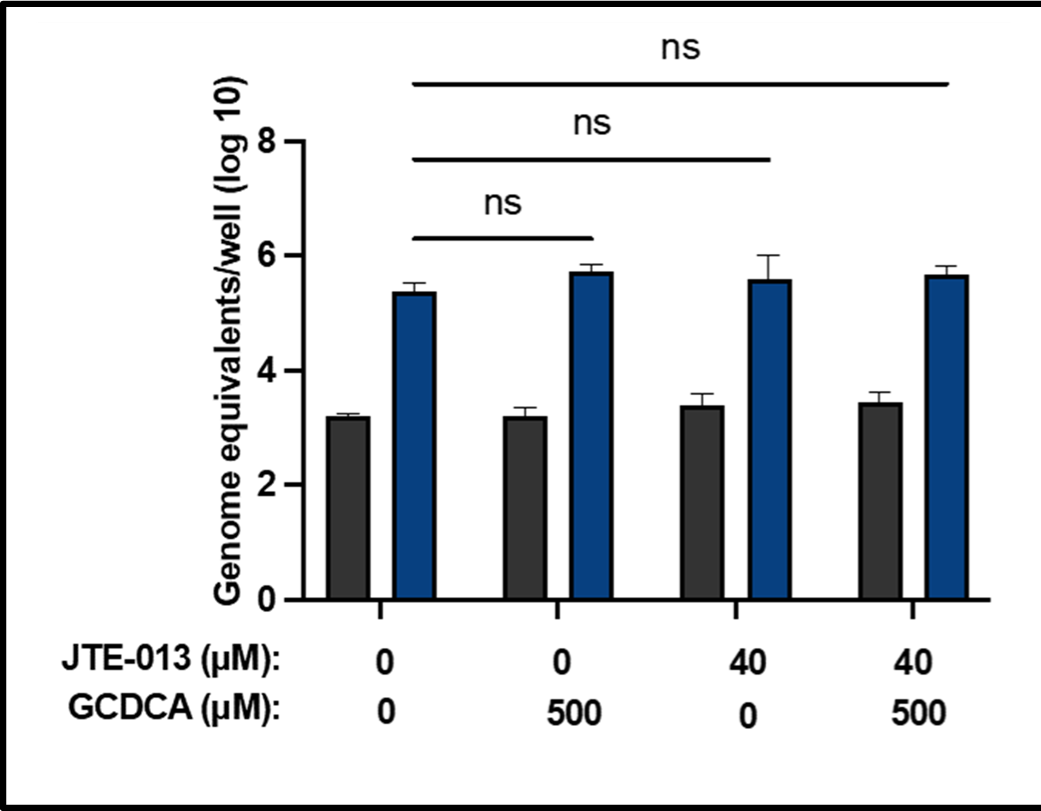
GII.4 replication is not reduced in the presence of JTE-013. GII.4 (4.3 × 10^5^ GEs/well) virus replication was evaluated in J2 HIEs plated in OGM in the presence or absence of GCDCA and JTE-013. The experiment was performed three times in triplicate, and a representative experiment is presented. Error bars denote standard deviation (n = 6). Significance was determined using two-way ANOVA.

We had previously shown that GII.4 Sydney enters HIEs by inducing apical ceramide appearance and LAMP-1 exocytosis and these molecules co-localize in the absence of BA treatment (14). We next tested whether S1PR2 was involved in these events in regard to GII.4 entry, as we showed above that BA treatment alone can induce apical ceramide generation and LAMP-1 exocytosis and their colocalization are affected by S1PR2 inhibition (**Figure 3**). We found that the GII.4 Sydney VLP-induced changes seen in the absence of exogenous BA treatment are not altered with S1PR2 inhibition (**Figure 5**).

**Figure 5.**
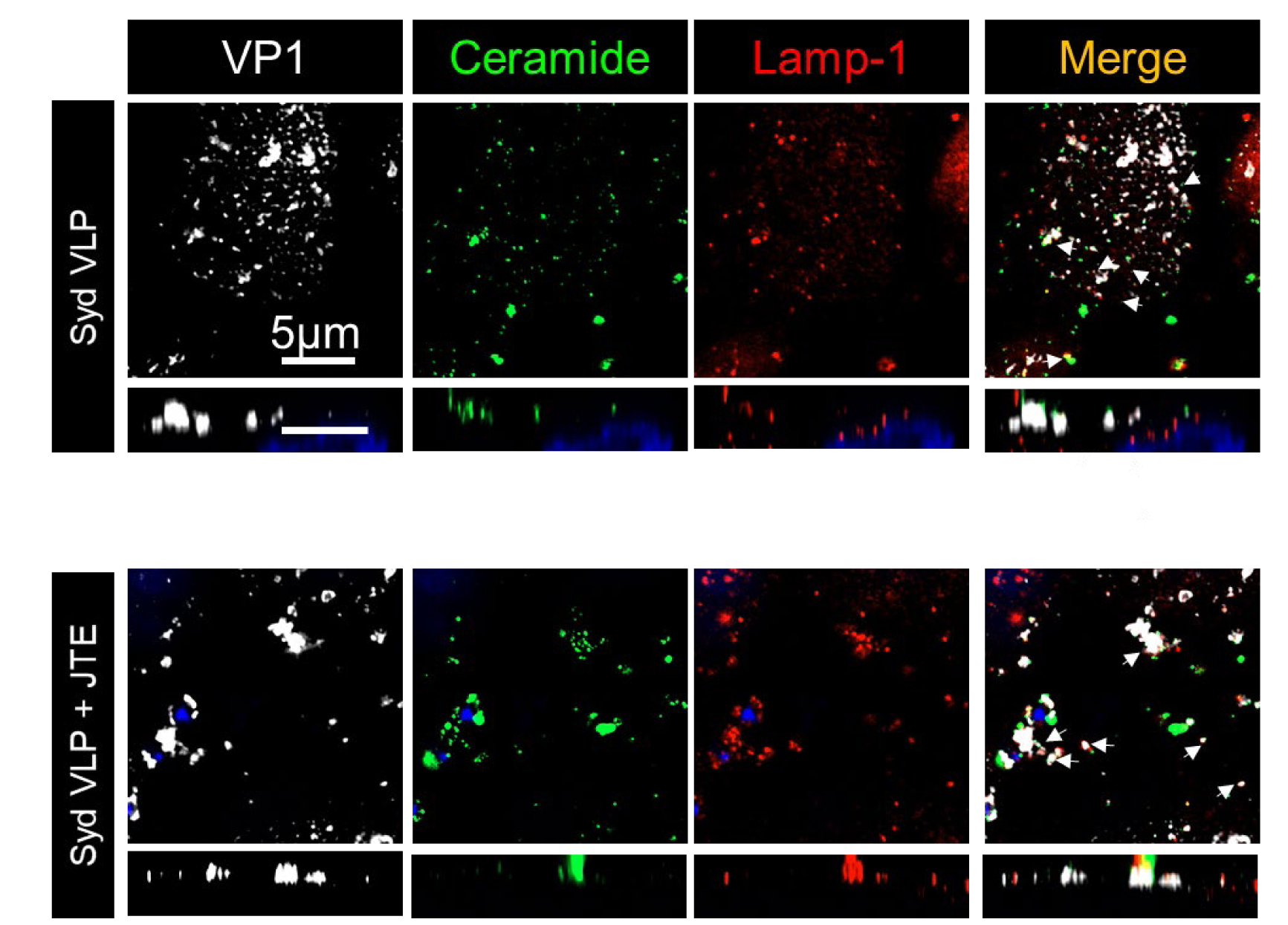
GII.4 Sydney VLP-induced apical ceramide appearance and LAMP-1 colocalization in HIEs are not regulated by S1PR2. Confocal microscopy shows detection of ceramide (green) and LAMP-1 (red) in J2 HIEs treated with GII.4 Sydney VLPs for 10 min in the absence (top panels) and presence (bottom panels) of the S1PR2 inhibitor, JTE-013. White arrows represent ceramide and LAMP-1 colocalization in JTE-013-treated vs untreated cells.

### Replication of other BA-dependent strains of HuNoV is also inhibited by JTE-013

We next sought to test other important BA-requiring strains of HuNoV. GII.17 HuNoVs caused local outbreaks such as an outbreak in a disaster relief megashelter in Houston, Texas following Hurricane Katrina (19). Similar to the GII.4 strain, variants of GII.17 have been proposed. Recently, a GII.17 variant maintained local predominance in Asia for multiple years (20–22). Two GII.17 isolates replicate in HIEs, and both require bile or BA (9, 13). Using the J4FUT2KI HIE line, where we previously showed that GII.17 replicates significantly better than in J2 (10), we tested the effect of S1PR2 inhibition. In the presence of JTE-013 during infection, GII.17 replication was significantly reduced compared to the control infection (**Figure 6A**). Next, we tested the effect of S1PR2 inhibition on a BA-requiring strain from the GI genogroup of HuNoVs. Replication of the prototype Norwalk strain, GI.1, was also reduced in the presence of JTE-013 during infection (**Figure 6B**). Taken together, these results indicate that sensitivity to S1PR2 inhibition is related to BA requirements and these strains likely share a common entry mechanism independent of their genogroup.

**Figure 6.**
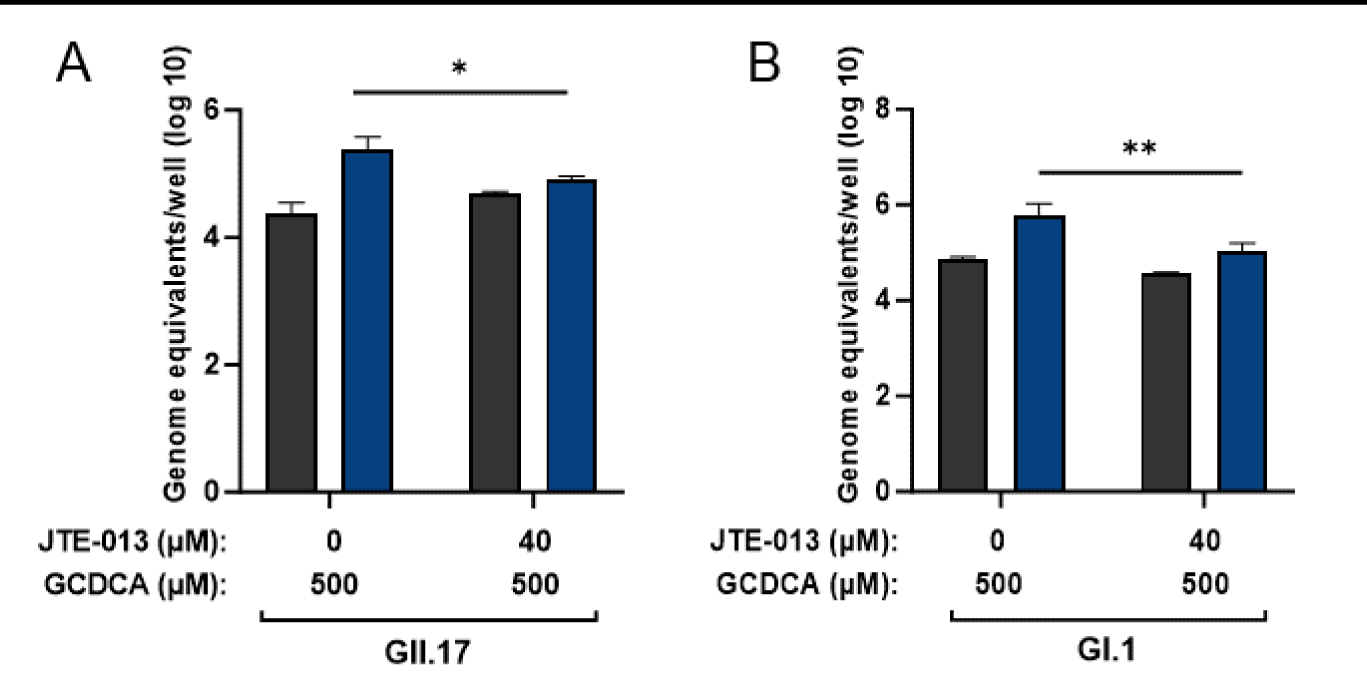
Replication of bile acid-dependent HuNoVs is reduced in the presence of JTE-013. (A) GII.17 (9.3 × 105 GEs/well) and (B) GI.1 (3.9 × 105 GEs/well) virus replication was evaluated in J4FUT2 KI HIEs plated in OGM and infected in the presence of GCDCA and in the presence or absence of JTE-013. Each experiment was performed twice, and compiled data are presented. Error bars denote standard deviation (n = 12). Significance was determined using a two-way ANOVA (P value -*, < 0.05; **, <0.01).

### BA-dependent replication of all segments of the small intestine requires S1PR2

Thus far our studies of BA-dependent HuNoV entry have focused on replication in jejunal HIEs, which led to the discovery that treatment of jejunal HIEs with GCDCA triggers endocytosis (11). However, GII.3 and other HuNoVs can infect other segments of the small intestine but not the colon (9, 13, 18). In the ileum, BAs are actively transported by the sodium-dependent bile acid transporter ASBT and interact with the nuclear receptor called farnesoid X receptor (FXR) (23). If the entry mechanism of GII.3 is similar in different segments of the small intestine, we anticipated S1PR2 inhibition would still have a strong negative effect on infection despite the higher presence of other BA-interacting receptors/transporters. Initially we used duodenal and ileal HIEs from one individual and observed that GII.3 infection was reduced in the presence of JTE-013 in both segments of the small intestine (not shown). Recently, we’ve added additional HIE lines to our bank including matched jejunal, duodenal, ileal, and colonic segments generated from the single individuals (designated 2004 and 2005). Infection results of the duodenal and jejunal HIE lines, D2004 and J2004, respectively, were similar to infection of the J2 line (**Figure 7**). The ileal line, IL2004, displayed over a log10 increase in genome equivalents at 24 hours post infection (hpi) compared to 1 hpi but did not reach levels as high as observed in the other segments. In all small intestinal segments from the 2004 individual, GII.3 infection was significantly decreased by the presence of JTE-013 during infection when compared to the untreated control infections. The colonic line, C2004 is not permissive to infection (Ettayebi et al., manuscript in preparation) and was not tested with JTE-013. Data from a set of HIEs from a different donor (2005) also showed GII.3 replication was inhibited by JTE-013 in the duodenum and jejunum from this individual but this virus did not grow in cultures from the ileum (**Figure 7**). This result demonstrates that the requirement for S1PR2 in BA-dependent GII.3 infection is conserved across infection of all segments of the small intestine indicating that the virus likely uses similar mechanisms for entry into cells in the duodenum, jejunum and ileum.

**Figure 7.**
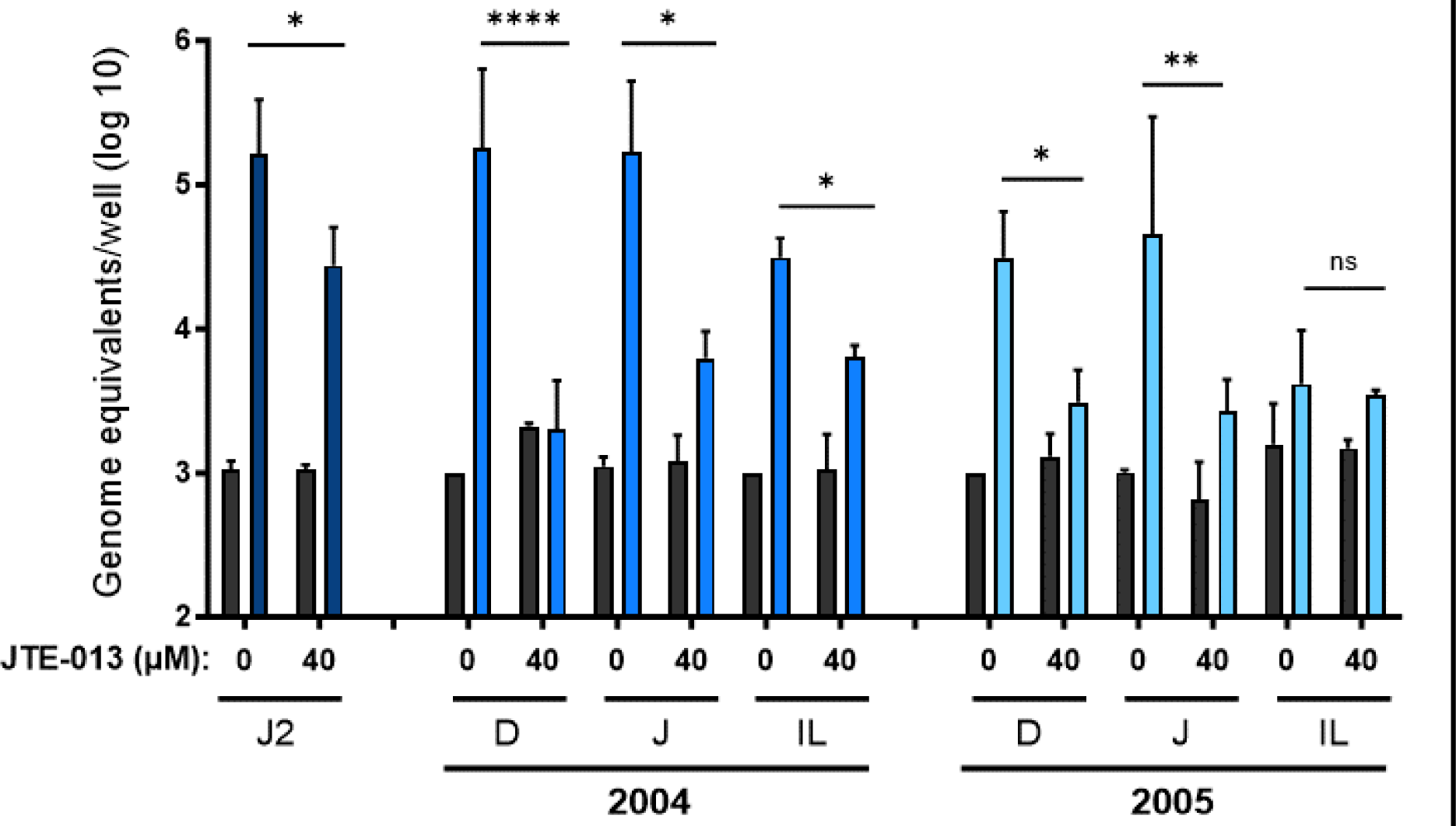
JTE-013 reduces the replication of GII.3 replication in HIEs from all segments of the small intestine. GII.3 (4.3 × 10^5^ GEs/well) virus replication was evaluated in J2 and HIEs derived from the duodenum (D), jejunum (J), and ileum (IL) from two individuals (2004 and 2005) plated in OGM and infected in the presence of GCDCA and absence or presence of JTE-013. Each experiment was performed twice, and compiled data are presented. Error bars denote standard deviation (n = 12). Significance was determined using a two-way ANOVA (P value - *, 0.05; **, 0.01; ****, < 0.001; ns, not significant).

## Discussion

The recent development of a small intestinal cultivation system has allowed us to begin probing differing requirements for the diverse strains of human norovirus replication. Our initial studies using jejunal HIEs demonstrate several key differences including that GII.3, but not GII.4, is sensitive to endogenous interferon; most strains tested except GII.4 require bile or BA for infection; and GII.3 requires BA to trigger cellular changes and S1PR2 for entry. In contrast, GII.4 will induce cell wounding and trigger CLIC entry pathways in the absence of BA (9, 11–14). In the current study, we confirmed the requirement for S1PR2 in GII.3 infection of a single jejunal HIE line and extended this conclusion to additional small intestinal lines and to other BA-dependent strains of HuNoV. Consistent with our observations that the pandemic GII.4 Sydney variant behaves differently from GII.3, infection by the pandemic strain was unaffected by the presence of the S1PR2 inhibitor, JTE-013. The different entry and infection requirements of the BA-independent GII.4 variants may be an explanation for why these are the predominant strains causing human disease. In addition to the effects of antagonizing S1PR2 in GII.3 infection, we found that BA treatment of HIEs rapidly enhances detection of this protein and the appearance of ceramide on the cell surface. These results suggest there are conformational changes in S1PR2 after BA addition likely associated with induced signaling pathways. These would be consistent with structural and functional properties of S1PR2 activation in other systems (24, 25). Finally, replication of two other BA-requiring strains, GII.17 and GI.1, was inhibited by S1PR2 inhibition, suggesting that a common entry and infection pathway is used by these strains independent of the genogroup (GI versus GII). When developing methods to mitigate infection by HuNoVs, it will be important to consider the effect on BA-independent GII.4s versus BA-requiring strains. Testing replication in the presence of BA with or without JTE-013 should be a useful new tool to classify HuNoVs into BA-dependent and -independent subgroups. Additionally, to achieve a higher level of GII.17 infection, sufficient to measure significant responses to JTE-013 treatment, we took advantage of the genetically-modified line (J4FUTKI) developed previously (10). Use of genetically-modified HIEs and culture conditions that promote the highest levels of infection will allow future studies on the infection mechanisms of less robust replicating genotypes of HuNoV.

In order to combat human disease including viral infection, drug development is important but extremely costly and has a high failure rate (26). One limitation is that laboratory cell culture lines do not represent the diversity of responses to a specific drug among patients and are often not the cell type the target virus infects (26–28). Use of complex multicellular cultures derived from multiple patients, such as with our HIE system, offers the opportunity to overcome these challenges early in the development pipeline when identifying potential drug targets (29, 30). HIEs recapitulate the intestinal epithelia and contain small intestinal enterocytes, accurately representing the tropism of HuNoV infection and we have derived our HIEs from many different patients and segments (31, 32). We have already used our expansive biobank of HIE lines as well as genetically-modified lines to evaluate a susceptibility factor for infection (HBGA) and developed a pipeline for testing antivirals (10, 18, 33). In this study, we used several different HIE lines generated from individuals that express secretor HBGAs and are permissive to infection by multiple strains of HuNoV, allowing us to explore the inhibitory effect of JTE-013 in infection of multiple lines. GII.3 replication was reduced by the S1PR2 inhibitor in several jejunal lines (J2, J3, J11, J4FUT2KI, J2004, and J2005). Moreover, factors that influence pharmacological efficacy such as absorption can vary throughout the small intestine. Here we show that GII.3 replication in genetically matched duodenal, jejunal, and ileal HIEs was inhibited through the use of JTE-013. Taken together, these studies highlight how future preclinical *in vitro* antiviral studies could utilize this system to compare the efficacy of drugs amongst multiple patients and in different small intestinal segments.

BAs have important functions in cholesterol homeostasis and to emulsify lipids during digestion processes but they also act as signaling molecules (34). Two well-studied receptors are the nuclear receptor FXR and the GPCR TGR5 (15). In our initial studies on the mechanism of BA-mediated GII.3 entry and the role of S1PR2, we used our main jejunal HIE line, J2, and drug modulation of FXR and TGR5 had no effect on infection (11). The major site of BA uptake is in the ileum via the apical sodium-dependent bile acid transporter (ASBT). Through transport into the cell, BAs gain access to and activate FXR in the ileum. TGR5 is also well expressed in the ileum where BA interactions with FXR and TGR5 can exert metabolic effects through signaling (35). In this study, we show that the less studied BA receptor S1PR2 is still an important BA receptor in GII.3 infection in the ileal HIE line derived from the patient 2004 despite the potential roles for FXR and TGR5 in the ileum. However, replication in the ileum was lower than in the matched jejunal and duodenal lines and absent in 2005. Future inhibitor experiments using permissive ileal HIE lines and agonists/antagonists of ASBT, FXR, and TGR5 will be necessary to understand whether these receptors also play a role in GII.3 replication in the ileum.

The natural ligand for S1PR2 is S1P but more recently in liver cells conjugated BAs were shown to activate downstream ERK1/2 and AKT signaling and modulate the activity of sphingosine kinase 2 (15, 16, 36). Knowledge of the role of BA signaling through S1PR2 in the intestine is limited. In intestinal epithelial cells, BA activation of S1PR2 had pro-proliferation effects; and in a mouse DSS model of colitis, the BA DCA activates cathepsin B release via S1PR2-ERK signaling and worsens the colitis (17, 37). We have now shown that S1PR2 inhibition blocks infection by BA-requiring strains of HuNoV and jejunal endocytosis (11). However, future studies are required to answer lingering questions and to further delineate the role S1PR2 plays in the small intestine and HuNoV infection: 1) Will S1P or agonists of S1PR2 have infection-promoting effects? 2) The lipid ceramide is important in HuNoV infection and a precursor to sphingosine, which is converted to S1P through sphingosine kinases. Will sphingosine kinase inhibitors alter HuNoV infection? 3) Recently, a group showed that JTE-013 has off-target effects inhibiting sphingosine kinases and dihydroceramide desaturase 1 (38). Will measurements of HIE cellular lipids after JTE-013 treatment reveal lipid changes that may affect infection? 4) Is ERK1/2 or AKT signaling downstream of S1PR2 activation important in mediating cellular changes required for HuNoV infection. 5) Finally, since our work has utilized exclusively inhibitor data, will knockout of S1PR2 in HIE lines prevent HuNoV infection?

In conclusion, we have shown that BAs increase detection of cellular S1PR2 and that colocalization of ceramide and LAMP-1 at the cell surface after BA treatment is blocked by inhibition of S1PR2. We have shown that BA-dependent HuNoV strains GII.3, GII.17 and GI.1 but not a BA-independent strain, GII.4 Sydney, require S1PR2 for infection. Using GII.3, these results were confirmed in HIE lines from several jejunal donors as well as in matched jejunal/duodenal/ileal sets of HIE lines from two individual donors. These results indicate the potential for targeting S1PR2 or subsequently activated pathways for therapeutic development. Future confirmatory studies on the role of S1PR2 and the JTE- 013 inhibitor in HuNoV infection will help further our understanding of these important gastrointestinal viruses and help elucidate additional roles that BAs and lipids play in intestinal biology.

## Methods

### Virus filtrates

Ten to 25% percent stool filtrates containing HuNoV were prepared as described previously (9). Briefly, 4.5 ml of ice-cold phosphate-buffered saline (PBS) was added to 0.5 ml of stool, homogenized by vortexing, and sonicated three times for 1 min. The sonicated suspension was centrifuged at 1,500 × g for 10 min at 4°C. The supernatant was transferred to a new tube and centrifuged a second time. The resulting supernatant was passed serially through 5-μm, 1.2-μm, 0.8-μm, 0.45-μm, and 0.22-μm filters depending on stool texture. The filtered sample was frozen in aliquots at −80°C until used. Virus stool filtrates are described in Table 1.

**Table 1.**

| Reference strain | Genotype | Filtrate<br>Percentage | Titer [G.E./μl] | G.E. per well<br>for infections |
| --- | --- | --- | --- | --- |
| GI.1[P1]/1968/Norwalk | GI.1 | 25% | $7.8 \times 10^4$ | $3.9 \times 10^5$ |
| GII.3[P21]/TCH04-577 | GII.3 | 10% | $8.5 \times 10^6$ | $4.3 \times 10^5$ |
| GII.4 Sydney[P31]/<br>TCH12-580 | GII.4 | 10% | $1.8 \times 10^7$ | $9.0 \times 10^5$ |
| GII.17[P13]/1295-44 | GII.17 | 10% | $9.3 \times 10^6$ | $9.3 \times 10^5$ |

### HIE lines

All HIE cultures used in this study are from an HIE bank maintained by the Texas Medical Center Digestive Diseases Center (TMC DDC) Core. Jejunal HIE cultures were previously generated from surgical specimens obtained during bariatric surgery. Duodenal, jejunal and ileal HIE cultures derived from the same individual (2204 or 2005) were obtained through the LifeGift tissue donation program. The BCM Institutional Review Board approved the study protocols. All HIEs used in this study were determined to be secretor-positive as previously described (9, 10). HIE cultures were grown as multilobular 3-dimensional (3D) HIEs in Matrigel and maintained in L-WRN complete media as described (13). For HuNoV infection, the 3D HIEs were dissociated by trypsinization and pipetting and seeded onto collagen IV coated 96-well plates. Monolayer cultures were seeded into 96-well plates using a 1:1 ratio of IntestiCult™ Organoid Growth Medium Human Basal Medium (StemCell Technologies, 100-0190) and Organoid Supplement (StemCell Technologies, 100-0191) (OGM proliferation medium) supplemented with 10 µM ROCK inhibitor Y-27632 for 24 hours. The HIEs were then differentiated for 5 days using a 1:1 ratio of Intesticult™ OGM and complete medium without growth factors (CMGF-, advanced DMEM/F12 prepared with 1X GlutaMAX and 10 mM HEPES) (OGM differentiation medium) with media changes every two days prior to inoculation with HuNoV.

### HuNoV Infections with JTE-013

A 50 mM stock solution of JTE-013 (Cayman) was prepared in fresh DMSO and frozen in single use aliquots used no longer than three months after preparation. Monolayer HIEs were pre-treated with 40 µM JTE-013 or vehicle control in OGM differentiation media for 1 h. Monolayers were inoculated in media with vehicle control or JTE-013 and with stool filtrate containing G.E.s of HuNoV per well as indicated in **Table 1** for 1 to 2 h (binding time increased to 2 h in some experiments as the longer incubation can increase replication levels) at 37°C in 5% CO2. Monolayers were then washed twice with CMGF(-) to remove unattached virus. The inoculated cells were maintained in OGM differentiation media with vehicle control or JTE-013 to 24 hpi.

### RNA extractions and RT-qPCR

Viral RNA was extracted at 1 or 2 h and 24 h using the KingFisher Flex purification system and the MagMAX-96 viral RNA isolation kit. G.E.s were determined by RT-qPCR as previously described (13). Briefly, the COG2R/QNIF2d/QNIFS (39) primer pair and probe were used for GII genotypes, and NIFG1F/NV1LCR/NIFG1P (40) were used for GI.1 with the qScript XLT One-Step RT-qPCR ToughMix reagent with ROX reference dye (Quanta Biosciences). Reactions were performed on an Applied Biosystems StepOnePlus thermocycler. A standard curve based on a recombinant HuNoV GII.4 (Houston virus) or GI.1 (Norwalk virus) RNA transcript was used to quantitate viral G.E.s in RNA samples (41, 42). Samples with RNA levels below the limit of detection of the RT-qPCR assay were assigned a value that was one-half the limit of detection of the assay.

### Western blotting

5-day differentiated J2 HIEs were incubated with 500 μM GCDCA for 10 min at 37°C in OGM differentiation media. The cells were washed twice and collected using a cell scrapper. HIEs without GCDCA treatment were used as a reference. S1PR2 expression in J2 HIEs was detected by Western blot analysis using rabbit S1PR2 polyclonal antibody (1:500; #AP01198PU-N, OriGene Technologies). Villin was used as cell loading control and detected using 1:1000 dilution of mouse anti-villin (#sc-373997, Santa Cruz Biotechnology) antibody.

### Confocal microscopy

Differentiated J2 HIEs were incubated with or without 500 μM GCDCA and 3 μg of GII.4 Sydney virus-like particles (VLPs) for 10 min at 37°C in OGM differentiation media (14). For testing the effect of inhibition of S1PR2 on GCDCA- or VLP- induced lysosomal exocytosis, the cells were preincubated with 40 μM of JTE-013 for 1 h prior to GCDCA or VLP addition and the inhibitor was also added during incubation of the cells with GCDCA and the VLPs. The cells were washed twice and fixed with 4% paraformaldehyde (PFA) at room temperature for 20 min. Cells were permeabilized with PBS containing 0.1% Triton x-100 and incubated overnight with primary antibodies (primary antibodies,1:200) at 4°C. S1PR2, ceramide, LAMP-1 and GII.4 VP1 were detected using rabbit S1PR2 polyclonal antibody, rabbit anti-ceramide polyclonal antibody (43), mouse anti-LAMP-1 monoclonal antibody (# sc-17768, Santa Cruz Biotechnology) and guinea pig anti-Sydney polyclonal antibody. After washing three times, the cells were incubated with phalloidin (1:1000, Invitrogen) for actin staining or a 1:500 dilution of secondary labeled antibody for 2 h at room temperature. The secondary antibodies used were [donkey anti-mouse 549 (#610-742-124, Rockland), donkey anti-rabbit 649 (#611-743-127, Rockland), and anti-guinea pig 488 (#606-141-129, Rockland)].

### Statistical Analysis

All statistical analyses were performed on GraphPad Prism for Windows (GraphPad Software, La Jolla, CA, USA). Each experiment was performed at least twice with three technical replicates in each condition. RT-qPCR assays were carried out using two or more technical replicates for each HIE replicate depending on the treatment of cells (represented by n). Data from compiled experiments or one representative experiment are presented. Comparison between 1 h and 24 h groups was performed using the Student’s t test, with statistical significance determined using the Holm–Sidak method. Where comparisons were made at 24 h between treatment groups, a two-way ANOVA was performed using Dunnett’s test for post hoc analyses. P <0.05 was considered statistically significant. Error bars denote standard deviation (SD).

## Acknowledgments

This research was supported in part by Public Health Service grants from the National Institutes of Health for grants P01 AI 057788 (to MKE), T32 AI055413 (to V.R.T.), S10 OD028480 that supported purchasing the Zeiss Laser Scanning Microscope LSM 980 with Airyscan 2, and P30 DK 056338 (to H. El-Serag), which supports the Texas Medical Center Digestive Diseases Center. The authors would like to acknowledge the Advanced Technology Core Laboratories (Baylor College of Medicine), specifically the Integrated Microscopy Core with funding from the NIH (DK56338, CA125123, ES030285), and CPRIT (RP150578, RP170719).

## References

1. Bagci S, Eis-Hubinger AM, Yassin AF, Simon A, Bartmann P, Franz AR, Mueller A. 2010. Clinical characteristics of viral intestinal infection in preterm and term neonates. Eur J Clin Microbiol Infect Dis 29:1079–84.

2. Brown LK, Clark I, Brown JR, Breuer J, Lowe DM. 2017. Norovirus infection in primary immune deficiency. Rev Med Virol 27:e1926.

3. Trivedi TK, Desai R, Hall AJ, Patel M, Parashar UD, Lopman BA. 2013. Clinical characteristics of norovirus-associated deaths: a systematic literature review. Am J Infect Control 41:654–7.

4. Koo HL, Neill FH, Estes MK, Munoz FM, Cameron A, DuPont HL, Atmar RL. 2013. Noroviruses: The most common pediatric viral enteric pathogen at a large university hospital after introduction of rotavirus vaccination. J Pediatric Infect Dis Soc 2:57-60.

5. Davis A, Cortez V, Grodzki M, Dallas R, Ferrolino J, Freiden P, Maron G, Hakim H, Hayden RT, Tang L, Huys A, Kolawole AO, Wobus CE, Jones MK, Karst SM, Schultz-Cherry S. 2020. Infectious Norovirus Is Chronically Shed by Immunocompromised Pediatric Hosts. Viruses 12.

6. Pires SM, Fischer-Walker CL, Lanata CF, Devleesschauwer B, Hall AJ, Kirk MD, Duarte AS, Black RE, Angulo FJ. 2015. Aetiology-Specific Estimates of the Global and Regional Incidence and Mortality of Diarrhoeal Diseases Commonly Transmitted through Food. PLoS One 10:e0142927.

7. Bartsch SM, Lopman BA, Ozawa S, Hall AJ, Lee BY. 2016. Global Economic Burden of Norovirus Gastroenteritis. PLoS One 11:e0151219.

8. Chhabra P, de Graaf M, Parra GI, Chan MC, Green K, Martella V, Wang Q, White PA, Katayama K, Vennema H, Koopmans MPG, Vinje J. 2019. Updated classification of norovirus genogroups and genotypes. J Gen Virol 100:1393–1406.

9. Ettayebi K, Crawford SE, Murakami K, Broughman JR, Karandikar U, Tenge VR, Neill FH, Blutt SE, Zeng XL, Qu L, Kou B, Opekun AR, Burrin D, Graham DY, Ramani S, Atmar RL, Estes MK. 2016. Replication of human noroviruses in stem cell-derived human enteroids. Science 353:1387–1393.

10. Haga K, Ettayebi K, Tenge VR, Karandikar UC, Lewis MA, Lin SC, Neill FH, Ayyar BV, Zeng XL, Larson G, Ramani S, Atmar RL, Estes MK. 2020. Genetic Manipulation of Human Intestinal Enteroids Demonstrates the Necessity of a Functional Fucosyltransferase 2 Gene for Secretor-Dependent Human Norovirus Infection. mBio 11.

11. Murakami K, Tenge VR, Karandikar UC, Lin SC, Ramani S, Ettayebi K, Crawford SE, Zeng XL, Neill FH, Ayyar BV, Katayama K, Graham DY, Bieberich E, Atmar RL, Estes MK. 2020. Bile acids and ceramide overcome the entry restriction for GII.3 human norovirus replication in human intestinal enteroids. Proc Natl Acad Sci U S A 117:1700-1710.

12. Lin SC, Qu L, Ettayebi K, Crawford SE, Blutt SE, Robertson MJ, Zeng XL, Tenge VR, Ayyar BV, Karandikar UC, Yu X, Coarfa C, Atmar RL, Ramani S, Estes MK. 2020. Human norovirus exhibits strain-specific sensitivity to host interferon pathways in human intestinal enteroids. Proc Natl Acad Sci U S A 117:23782–23793.

13. Ettayebi K, Tenge VR, Cortes-Penfield NW, Crawford SE, Neill FH, Zeng XL, Yu X, Ayyar BV, Burrin D, Ramani S, Atmar RL, Estes MK. 2021. New Insights and Enhanced Human Norovirus Cultivation in Human Intestinal Enteroids. mSphere 6.

14. Ayyar BV, Ettayebi K, Salmen W, Karandikar UC, Neill FH, Tenge VR, Crawford SE, Bieberich E, Prasad BVV, Atmar RL, Estes MK. 2023. CLIC and membrane wound repair pathways enable pandemic norovirus entry and infection. Nat Commun 14:1148.

15. Ticho AL, Malhotra P, Dudeja PK, Gill RK, Alrefai WA. 2019. Bile Acid Receptors and Gastrointestinal Functions. Liver Res 3:31–39.

16. Nagahashi M, Takabe K, Liu R, Peng K, Wang X, Wang Y, Hait NC, Wang X, Allegood JC, Yamada A, Aoyagi T, Liang J, Pandak WM, Spiegel S, Hylemon PB, Zhou H. 2015. Conjugated bile acid-activated S1P receptor 2 is a key regulator of sphingosine kinase 2 and hepatic gene expression. Hepatology 61:1216–26.

17. Chen T, Huang Z, Liu R, Yang J, Hylemon PB, Zhou H. 2017. Sphingosine-1 phosphate promotes intestinal epithelial cell proliferation via S1PR2. Front Biosci (Landmark Ed) 22:596–608.

18. Lewis MA, Cortes-Penfield NW, Ettayebi K, Patil K, Kaur G, Neill FH, Atmar RL, Ramani S, Estes MK. 2023. Standardization of an antiviral pipeline for human norovirus in human intestinal enteroids demonstrates nitazoxanide has no to weak antiviral activity. Antimicrob Agents Chemother 67:e0063623.

19. Yee EL, Palacio H, Atmar RL, Shah U, Kilborn C, Faul M, Gavagan TE, Feigin RD, Versalovic J, Neill FH, Panlilio AL, Miller M, Spahr J, Glass RI. 2007. Widespread outbreak of norovirus gastroenteritis among evacuees of Hurricane Katrina residing in a large “megashelter” in Houston, Texas: lessons learned for prevention. Clin Infect Dis 44:1032–9.

20. Jin M, Zhou YK, Xie HP, Fu JG, He YQ, Zhang S, Jing HB, Kong XY, Sun XM, Li HY, Zhang Q, Li K, Zhang YJ, Zhou DQ, Xing WJ, Liao QH, Liu N, Yu HJ, Jiang X, Tan M, Duan ZJ. 2016. Characterization of the new GII.17 norovirus variant that emerged recently as the predominant strain in China. J Gen Virol 97:2620- 2632.

21. Chan MC, Lee N, Hung TN, Kwok K, Cheung K, Tin EK, Lai RW, Nelson EA, Leung TF, Chan PK. 2015. Rapid emergence and predominance of a broadly recognizing and fast-evolving norovirus GII.17 variant in late 2014. Nat Commun 6:10061.

22. de Graaf M, van Beek J, Vennema H, Podkolzin AT, Hewitt J, Bucardo F, Templeton K, Mans J, Nordgren J, Reuter G, Lynch M, Rasmussen LD, Iritani N, Chan MC, Martella V, Ambert-Balay K, Vinje J, White PA, Koopmans MP. 2015. Emergence of a novel GII.17 norovirus - End of the GII.4 era? Euro Surveill 20.

23. Chiang JY. 2013. Bile acid metabolism and signaling. Compr Physiol 3:1191–212.

24. Andermatten RB, Ciriaci N, Schuck VS, Di Siervi N, Razori MV, Miszczuk GS, Medeot AC, Davio CA, Crocenzi FA, Roma MG, Barosso IR, Sanchez Pozzi EJ. 2019. Sphingosine 1-phosphate receptor 2/adenylyl cyclase/protein kinase A pathway is involved in taurolithocholate-induced internalization of Abcc2 in rats. Arch Toxicol 93:2279–2294.

25. Chen H, Chen K, Huang W, Staudt LM, Cyster JG, Li X. 2022. Structure of S1PR2-heterotrimeric G(13) signaling complex. Sci Adv 8:eabn0067.

26. Sun D, Gao W, Hu H, Zhou S. 2022. Why 90% of clinical drug development fails and how to improve it? Acta Pharm Sin B 12:3049–3062.

27. Andrei G. 2021. Vaccines and Antivirals: Grand Challenges and Great Opportunities. Frontiers in Virology 1.

28. Eastman A. 2017. Improving anticancer drug development begins with cell culture: misinformation perpetrated by the misuse of cytotoxicity assays. Oncotarget 8:8854–8866.

29. Langhans SA. 2018. Three-Dimensional in Vitro Cell Culture Models in Drug Discovery and Drug Repositioning. Front Pharmacol 9:6.

30. Artegiani B, Clevers H. 2018. Use and application of 3D-organoid technology. Hum Mol Genet 27:R99–R107.

31. Karandikar UC, Crawford SE, Ajami NJ, Murakami K, Kou B, Ettayebi K, Papanicolaou GA, Jongwutiwes U, Perales MA, Shia J, Mercer D, Finegold MJ, Vinje J, Atmar RL, Estes MK. 2016. Detection of human norovirus in intestinal biopsies from immunocompromised transplant patients. J Gen Virol 97:2291–2300.

32. Ramani S, Crawford SE, Blutt SE, Estes MK. 2018. Human organoid cultures: transformative new tools for human virus studies. Curr Opin Virol 29:79–86.

33. Tenge VR, Hu L, Prasad BVV, Larson G, Atmar RL, Estes MK, Ramani S. 2021. Glycan Recognition in Human Norovirus Infections. Viruses 13.

34. Thibaut MM, Bindels LB. 2022. Crosstalk between bile acid-activated receptors and microbiome in entero-hepatic inflammation. Trends Mol Med 28:223–236.

35. Chiang JY. 2017. Recent advances in understanding bile acid homeostasis. F1000Res 6:2029.

36. Nagahashi M, Yuza K, Hirose Y, Nakajima M, Ramanathan R, Hait NC, Hylemon PB, Zhou H, Takabe K, Wakai T. 2016. The roles of bile acids and sphingosine- 1-phosphate signaling in the hepatobiliary diseases. J Lipid Res 57:1636–43.

37. Zhao S, Gong Z, Du X, Tian C, Wang L, Zhou J, Xu C, Chen Y, Cai W, Wu J. 2018. Deoxycholic Acid-Mediated Sphingosine-1-Phosphate Receptor 2 Signaling Exacerbates DSS-Induced Colitis through Promoting Cathepsin B Release. J Immunol Res 2018:2481418.

38. Pitman MR, Lewis AC, Davies LT, Moretti PAB, Anderson D, Creek DJ, Powell JA, Pitson SM. 2022. The sphingosine 1-phosphate receptor 2/4 antagonist JTE- 013 elicits off-target effects on sphingolipid metabolism. Sci Rep 12:454.

39. Loisy F, Atmar RL, Guillon P, Le Cann P, Pommepuy M, Le Guyader FS. 2005. Real-time RT-PCR for norovirus screening in shellfish. J Virol Methods 123:1–7.

40. Miura T, Parnaudeau S, Grodzki M, Okabe S, Atmar RL, Le Guyader FS. 2013. Environmental detection of genogroup I, II, and IV noroviruses by using a generic real-time reverse transcription-PCR assay. Appl Environ Microbiol 79:6585–92.

41. Guix S, Asanaka M, Katayama K, Crawford SE, Neill FH, Atmar RL, Estes MK. 2007. Norwalk virus RNA is infectious in mammalian cells. J Virol 81:12238–48.

42. Le Guyader FS, Le Saux JC, Ambert-Balay K, Krol J, Serais O, Parnaudeau S, Giraudon H, Delmas G, Pommepuy M, Pothier P, Atmar RL. 2008. Aichi virus, norovirus, astrovirus, enterovirus, and rotavirus involved in clinical cases from a French oyster-related gastroenteritis outbreak. J Clin Microbiol 46:4011–7.

43. Krishnamurthy K, Dasgupta S, Bieberich E. 2007. Development and characterization of a novel anti-ceramide antibody. J Lipid Res 48:968–75.

